# Terminal flourishes but not trills differ between urban and rural chaffinch song

**DOI:** 10.1101/2022.06.29.498165

**Authors:** Alper Yelimlieş, Berkay Atalas, Çağla Önsal, Çağlar Akçay

## Abstract

Anthropogenic noise interrupts the acoustic communication between animals living in urban habitats. Accumulating evidence suggests that animals can evade this interruption using various strategies such as shifting frequencies upwards or increasing the duration of their signals. In this study, we compared the time and frequency-related characteristics of songs and rain calls of common chaffinches (*Fringilla coelebs*) inhabiting rural forests and an urban park in Turkey. Most of the song phrases and rain calls did not differ in any of the characteristics measured between urban and rural chaffinches. Terminal flourish phrases of the songs, however, had lower minimum frequencies and broader bandwidth in urban territories, contrary to our predictions. We discuss this finding in relation to its potential adaptive significance.

---

Acoustic communication between animals is constrained by abiotic and biotic sources of external noise such as wind, rainfall, streams, and vocalizations of other animals (Endler, 1992). Signals often evolve to adapt to the environmental conditions that they are to be transmitted (Hansen, 1979; Morton, 1975). While in natural habitats environmental change may be gradual, in a rapidly urbanizing world, communication channels are becoming increasingly polluted by anthropogenic noise (Brumm & Slabbekoorn, 2005). Consequently, the effect of noise on animal communication, particularly in the acoustic modality, has been a major focus of research (Halfwerk & Slabbekoorn, 2015).

Animals employ different strategies in adjusting their vocal communication to cope with noise (Brumm & Slabbekoorn, 2005; Kunc & Schmidt, 2021; Patricelli & Blickley, 2006). One widespread strategy among animals is simply increasing the amplitude of their signals (Brumm & Zollinger, 2011). Moreover, animals may alter the timing of their signals to avoid predictable noise (Arroyo-Solís et al., 2013), increase the redundancy of their acoustic signals by coupling them with signals in other modalities carrying the same information (Akçay & Beecher, 2019), or repeat the same signal more frequently in noisy conditions (Brumm & Slater, 2006; Kaiser & Hammers, 2009).

Animals also alter the acoustic structure of their vocalizations in response to noise, by changing temporal or frequency parameters, such as by increasing the duration or the minimum frequency of vocalizations to avoid masking caused by low-frequency anthropogenic noise (Ríos-Chelén et al., 2013; Slabbekoorn & Peet, 2003; Wood & Yezerinac, 2006). If a shift in minimum frequency is not accompanied by an equivalent increase in higher maximum frequency, the narrower bandwidth of the signal can also increase its transmission probability (Lohr et al., 2003; Winandy et al., 2021). Additionally, animals can diminish masking by shifting the energy concentration of their signals (i.e. peak frequency) to higher frequencies (Francis et al., 2011; Shieh et al., 2012).

Although the changes in acoustic structure may facilitate transmission of the signal in a noisy environment, they may also result in a trade-off with other fitness-relevant functions of vocalizations. In some species, where vocalizations with lower frequencies indicate the size or quality of a mate, evading masking by increasing the minimum frequency of a signal may come at the expense of reproductive success (Halfwerk, Bot, et al., 2011; Halfwerk, Holleman, et al., 2011). Also, as signals with higher frequency bandwidths are also important for demonstrating quality in intrasexual interactions, decreased bandwidth associated with a higher minimum frequency of a song for increasing transmission probability evokes less response in competitor males (Luther et al., 2016). Moreover, as urbanization alters many other characteristics of habitats, alterations in signals may be constrained by better functioning in other fitness-relevant activities such as bill morphology adapted to new foraging options affecting signal production (Giraudeau et al., 2014).

With its well-studied communication system, the common chaffinch (*Fringilla coelebs*, hereafter chaffinch) is a suitable species for asking questions about the influence of anthropogenic noise on vocalizations. The chaffinch is a territorial songbird and males are commonly found singing throughout urban and rural habitats in spring. They have a repertoire of two to six discrete song types which they sing with eventual variety (singing one type of song repeatedly before changing to another) with some types favored over others (Hinde, 1958; Slater, 1981). Each song has a structure that consists of a varying number of trill phrases with repeating syllables, and a *flourish*, which is a more complex end phrase containing different note types (Marler, 1956; see Figure 1).

**Figure 1.**
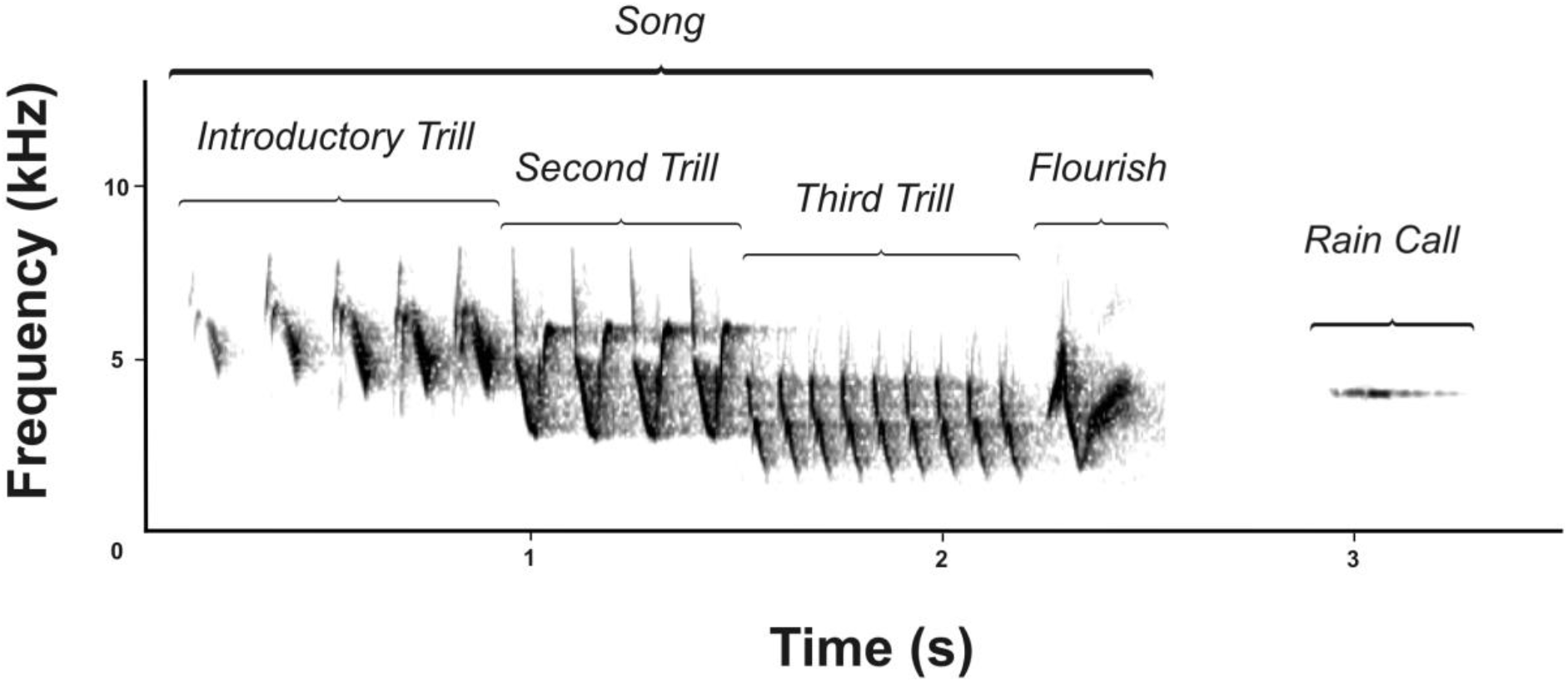
Spectrogram of chaffinch signals: a song with three trill phrases along with a terminal flourish phrase, and a rain call.

Different parts of the chaffinch song may carry different messages for various receivers. Early studies showed that the first two phrases of a song are the most important components for species recognition (Brémond, 1972). Observational studies suggest that the trill phrases may have a function in aggressive interactions between males: males sing incomplete songs (songs with trills but without the flourish) more often in an agonistic interaction with a conspecific, and increased numbers of incomplete songs often lead to fighting between males (Heymann & Bergmann, 1988). Playbacks of trills alone elicit a more aggressive response from males compared to flourish alone (Brémond, 1972). Finally, songs with longer trills evoke stronger responses from males than songs with shorter trills controlling for the song duration (Leitão & Riebel, 2003). All of these findings suggest that trill phrases are functional in male-male communication.

In contrast to trill phrases, flourish phrases may be a signal primarily directed at females. Evidence for this hypothesis comes from a study that found songs with flourishes, and among them, those with relatively longer flourishes were preferred more by female chaffinches as indicated by their perch selection which activates song playbacks (Riebel & Slater, 1998). Moreover, chaffinches tend to sing songs with repeating flourishes when they are sexually stimulated (Marler, 1956). Together these findings suggest that trills and flourishes may function in different contexts: the former mostly serving a male-male signaling function while the latter functioning mostly as a female-directed signal.

During the breeding season, male chaffinches also use the “rain calls” (so-called because of an apparent association with adverse weather) which appear to be used in multiple contexts although the precise function is unclear (Marler, 1956; Poulsen, 1958). Several authors describe rain calls given in situations where “song motivation is low or song is inhibited” (Poulsen, 1958, p. 93). Rain calls are also used as an alarm signal in response to predators, although as the intensity of the alarm increases rain calls are gradually replaced with the chink calls (Krams & Krama, 2002). In other contexts, rain calls are associated with courtship, within-pair communication, and parent-offspring communication (Marler, 1956). This call type geographically varies in its structure (Skiba, 2000; Sorjonen, 2001), which can be tonal (as in our population, see Figure 1) or resemble a trill or buzz (see Figure 3 in Skiba, 2000).

While many studies examined the function of chaffinch song and calls, few studies examined how these signals change with noise. In a seminal study, Brumm and Slater (2006) examined the singing behavior of chaffinches depending on naturally occurring noises such as waterfalls and strong streams. They found chaffinches increased the serial redundancy of their signal by repeating a song type longer than chaffinches in quieter habitats (Brumm & Slater, 2006). A more recent study, however, did not find the effect in a comparison of chaffinch songs between quiet rural and noisy urban areas (Deoniziak & Osiejuk, 2016). With respect to the acoustic features of the songs, Brumm and Slater (2006) reported that chaffinches increase the minimum frequency of their songs to reduce masking by low-frequency noise. Similarly, Kislyakov and Ivanitskii (2019) found that increased levels of anthropogenic noise were strongly associated with higher minimum frequency in 3 out of 4 types of trill phrases of 3 types of songs shared across chaffinches in their study area. In addition to the songs, Skiba (2000) investigated the rain call structure in relation to traffic noise but did not find any association between the distribution of different dialect types and acoustic characteristics of the environment. Moreover, Skiba (2000) found that spectral characteristics of the rain calls were suboptimal for their transmission in noise.

Here we compare acoustic features of chaffinch song phrases and rain calls between urban and rural areas. Considering the masking that would be caused by low-frequency noise in cities, we hypothesized that there would be a shift towards higher minimum frequencies in all signal types of urban living chaffinches. In addition, we expected them to be longer in duration in urban chaffinches compared to rural chaffinches. Moreover, we investigated the speed of trilled song parts to see if males compensate for a possible bandwidth loss with higher trill rates.

## Methods

### Study site and subjects

We recorded 28 male chaffinches between 11 and 26 May 2021 in Sariyer, Istanbul, Turkey. Rural individuals were inhabiting forests near Koç University’s main campus in the north of Istanbul, whereas urban ones were inhabiting a large urban park (Atatürk Kent Ormani, Haciosman, Sariyer: 41°08’05.4”N, 29°01’55.5”E). Out of those recordings, songs were obtained from 25 individuals (14 rural, 11 urban), and rain calls were obtained from 15 males (8 rural, 7 urban). All of the recorded birds were unbanded, so we selected territories that are at least 150 meters apart from each other for recordings in order to confidently assign separate identities to them. We did not attempt to record the entire repertoire of the males.

In focal territories, except for four of them, we measured ambient noise following the procedure described by Brumm (2004). We took two samples from each of the four perpendicular directions using a sound level meter (VLIKE VL6708, VLIKE Inc.), then we calculated their average. The four territories that do not have ambient noise measurements were situated in field sites that contain territories for which ambient noise was measured.

### Recordings

The number of songs obtained from each male ranged between 1 and 45 (*M* = 14.44, *SD* = 11.63) for a total of 361 songs. These recordings were collected during free singing (n = 26) or in response to brief playbacks of the conspecific song (n = 335). Whether or not the songs were recorded in response to playback did not have a significant effect on any of the measures below. All of the rain call recordings were obtained after broadcasting song playbacks. The number of them obtained from each individual varied between 8 and 309 (*M* = 104.67, *SD* = 80.28) for a total of 1570 calls. Taken together, we recorded 198 songs and 737 rain calls from rural chaffinches along with 163 songs and 833 rain calls from urban chaffinches. We used a Marantz PMD660 recorder with a Sennheiser ME66/K6 or ME67/K6 shotgun microphone for the recordings (sampling rate: 44100 Hz, 16 bit).

### Acoustic Analysis

Using Raven Pro 1.6.3 (K. Lisa Yang Center for Conservation Bioacoustics, 2022), we created spectrograms for recordings (Hann window, window length= 512, overlap= 50%). Then, we manually selected rain calls and song phrases from the spectrograms and saved these selections. We named phrases with respect to their type and the order it occurs in the song (i.e. introductory trill, 2^nd^ trill, 3^rd^ trill and flourish). The number of thrill phrases in a song varied in our sample with a maximum number of four, however fourth trill phrases occurred only in 50 songs, and among them, only 2 were from urban males. Considering its limited sample size, we removed fourth trill phrases from all of our analyses.

For the time and frequency-related parameters, we used methods that yield bias-free measurements as determining them visually (“eye-balling”) on the spectrogram is known to cause bias in minimum frequency measurements in noisy recordings and is therefore unreliable (Brumm et al., 2017). We therefore used the threshold method (i.e. setting an amplitude threshold relative to the peak in power spectra, see Figure 2, Zollinger et al., 2012) for measuring the bandwidth, minimum and maximum frequency by adapting the custom R code (R Core Team, 2021) written by Clink and colleagues (2018). We set the threshold at -12 dB for song phrases and -15 dB for the rain calls, as the latter signal type had energy concentration at higher frequencies relative to the noise compared to the former. We screened data and removed these measures from 71 song phrases as the minimum frequency indicated by this method seemed to be background noise. Among those phrases, 59 were from urban habitats, 51 were introductory trills (these tended to be sometimes relatively quiet), 16 were flourishes, 3 were second trills and 1 was a third trill.

**Figure 2.**
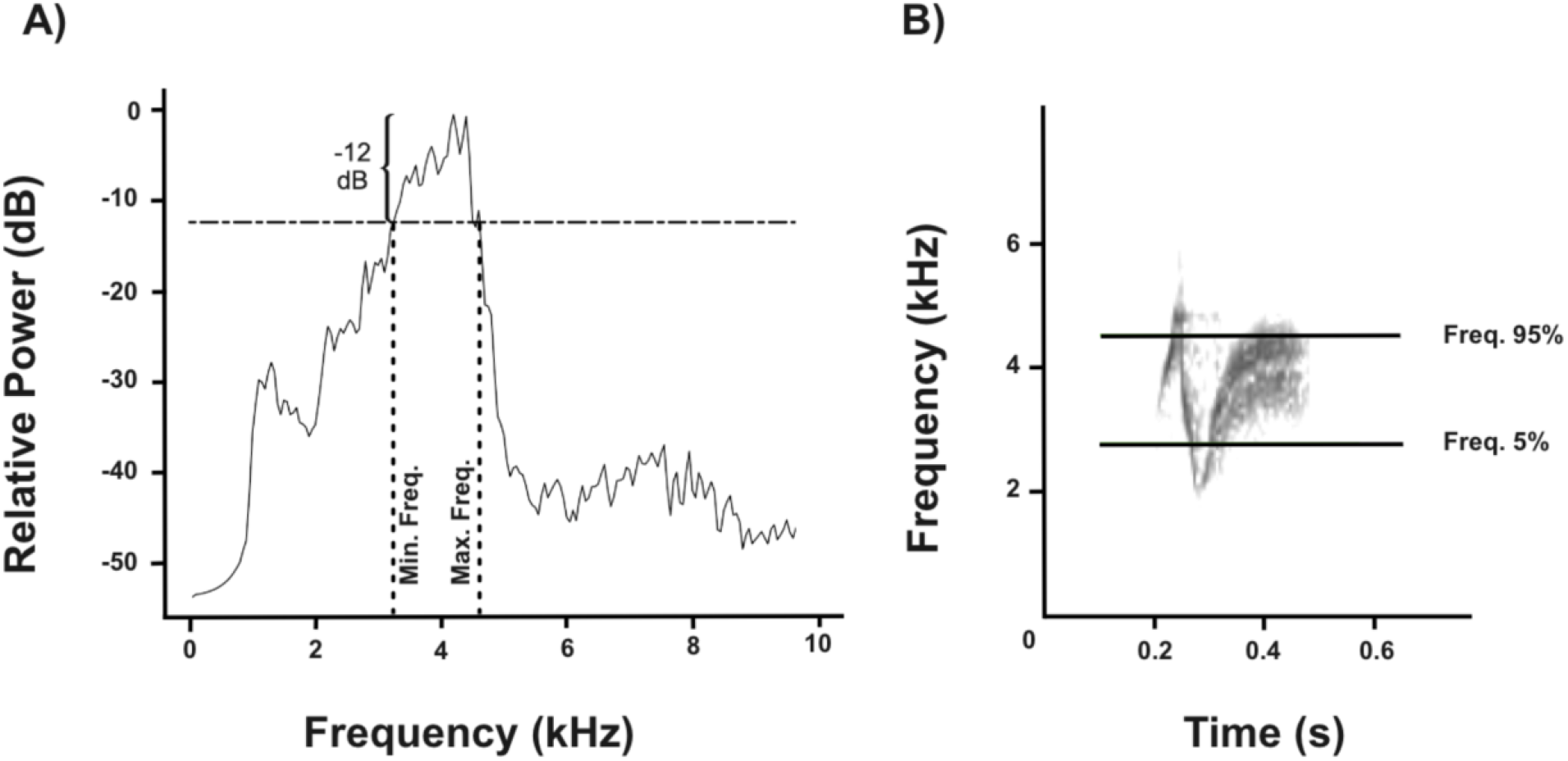
A) Power spectra illustrating the minimum and maximum frequency of a flourish phrase measured at -12 dB. B) Spectrogram of the same phrase showing energy distribution measurements marked by Raven Pro 1.6.3.

We used the built-in functions of Raven Pro for the rest of the measures. The *Peak frequency* gives the frequency value which has the maximum amplitude for each phrase. We also included energy distribution measures for frequency as it is not redundant with other measures and they do not get influenced by how selection borders were set around the signal (Charif et al., 2010). For the energy distribution of each phrase, we used *Frequency 5%, Frequency 95%*, and *Bandwidth 90%* which gives the frequency value that contains the 5% and 95% of the energy below it in the selection, and the difference between the first two measures, respectively. Considering the simple tonal structure of the rain calls, we did not include the energy distribution parameters for them as these were largely redundant with the minimum and maximum frequency measures made with the threshold method. Lastly, we measured the duration of each vocalization using the *Duration 90%* function which gives the time range that is between the 5^th^ and 95^th^ percentiles of the energy concentration in seconds. To quantify the trill rates, we counted the number of syllables in each selection and divided that number by the duration of the selection. In many utterances of the introductory trill, chaffinches started singing softly. As this weak signal-to-noise ratio in the beginning significantly influenced the measurement of duration, we did not quantify trill rates for introductory trills.

### Data analysis

To assess the differences between habitats in the acoustic parameters of chaffinch song, we constructed linear mixed models (LMMs) using the *nlme* package in R (Pinheiro et al., 2022) first to investigate ambient noise differences between habitats and then to test the spectral and temporal differences between song phrases from urban and rural habitats. Our model with the ambient noise level as the dependent variable had the habitat (rural vs. urban) as the fixed factor and bird ID as the random factor as some birds were recorded on two consecutive days. For every suitable parameter, we ran LMMs for each phrase type (introductory trill, second trill, third thrill and flourish) separately with habitat as the fixed factor and bird ID as the random factor. We also ran LMMs with the same fixed and random factors for rain call parameters.

## Results

Urban and rural habitats inhabited by chaffinches in our study significantly differed in the volume of their ambient noise. As expected, average ambient noise was louder in urban habitats compared to rural ones (LMM: coefficient ± SE: 9.30 ± 1.53, χ^2^ = 36.94, *P* = 0.000000001, marginal R^2^ = 0.59, see Figure 2).

Acoustic parameters of introductory, second, and third trill phrases did not differ between urban and rural habitats (Tables 1-3). Flourish phrases had lower minimum frequencies (LMM: coefficient ± SE: -385.29 ± 181.26, χ^2^ = 4.53, *P* = 0.03, marginal R^2^ =0.09) and broader bandwidths (LMM: coefficient ± SE: 500.09 ± 222.50, χ^2^ = 5.05, *P* =0.02, marginal R^2^ = 0.09; Table 5, see Figure 3) in urban habitats compared to rural habitats. Energy distribution measurements also paralleled this finding as flourish phrases had greater Bandwidth 90% values in urban songs (LMM: coefficient ± SE: 404.183 ± 189.52, χ^2^ = 4.56, *P* = 0.03 marginal R^2^ =0.08). The other parameters of flourish phrases did not differ significantly between urban and rural habitats (Table 4). Rain calls did not differ between the habitats in any of the acoustic measures (Table 5).

**Table 1.** Linear mixed models for introductory trill with the habitat type as the fixed factor (baseline = rural, n =310 for parameters measured with threshold method, for the rest n = 361)

| Introductory Trill Phrase |  |  |  |
| --- | --- | --- | --- |
| | Coefficient (SE) | $\chi^2$ | P-value |
| Min. Frequency | -46.69 (228.14) | 0.04 | 0.84 |
| Max. Frequency | 203.95 (188.10) | 1.18 | 0.28 |
| Bandwidth | 222.37 (289.35) | 0.59 | 0.44 |
| Frequency 5% | 7.55 (124.74) | 0.004 | 0.95 |
| Frequency 95% | 14.82 (152.12) | 0.01 | 0.92 |
| Bandwidth 90% | -3.21 (155.02) | 0.0004 | 0.98 |
| Peak Frequency | -65.11 (220.47) | 0.09 | 0.77 |
| Duration 90% | 0.06 (0.04) | 2.15 | 0.14 |

**Table 2.** Linear mixed models for the second trill with the habitat type as the fixed factor (baseline = rural, n = 358 for parameters measured with threshold method, for the rest n = 361)

| Second Trill Phrase |  |  |  |
| --- | --- | --- | --- |
| | Coefficient (SE) | $\chi^2$ | P-value |
| Min. Frequency | -68.73 (135.41) | 0.26 | 0.61 |
| Max. Frequency | 178.29 (324.79) | 0.30 | 0.58 |
| Bandwidth | 280.84 (241.75) | 1.35 | 0.25 |
| Frequency 5% | -106.10 (106.90) | 0.99 | 0.32 |
| Frequency 95% | 151.82 (226.23) | 0.45 | 0.50 |
| Bandwidth 90% | 278.40 (182.36) | 2.33 | 0.13 |
| Peak Frequency | -119.60 (159.35) | 0.56 | 0.45 |
| Duration 90% | 0.013 (0.06) | 0.05 | 0.83 |
| Trill Rate | -0.61 (1.41) | 0.19 | 0.66 |

**Table 3.**
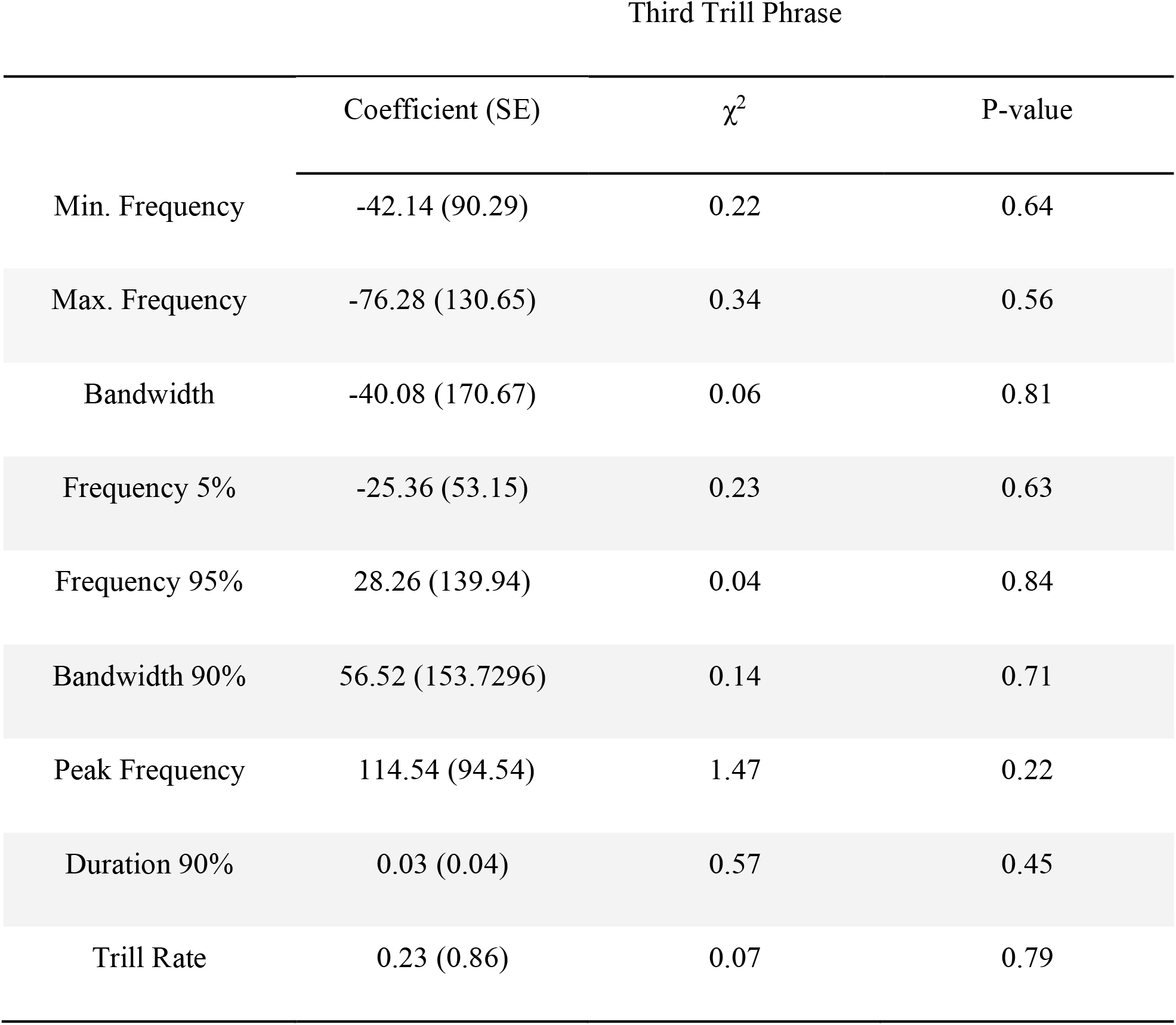
Linear mixed models for the third trill with the habitat type as the fixed factor (baseline = rural, n =323 for parameters measured with threshold method, for the rest n = 324)

**Table 4.**
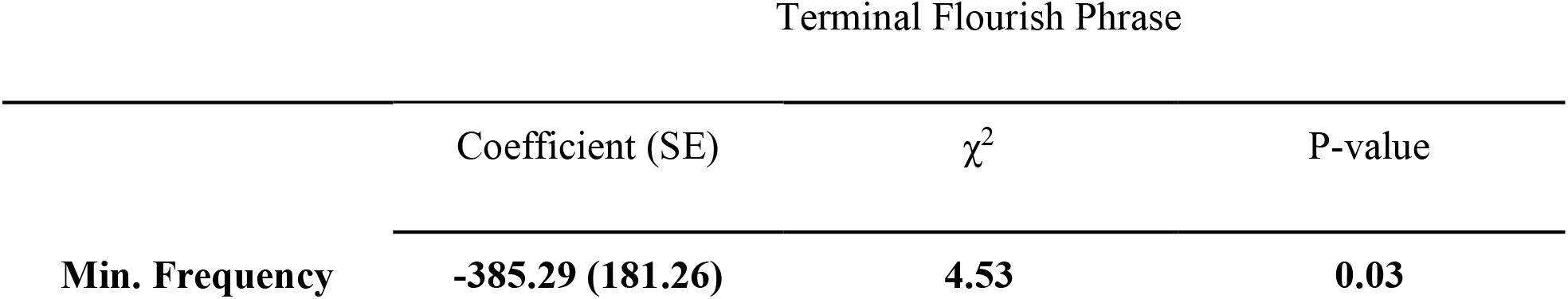
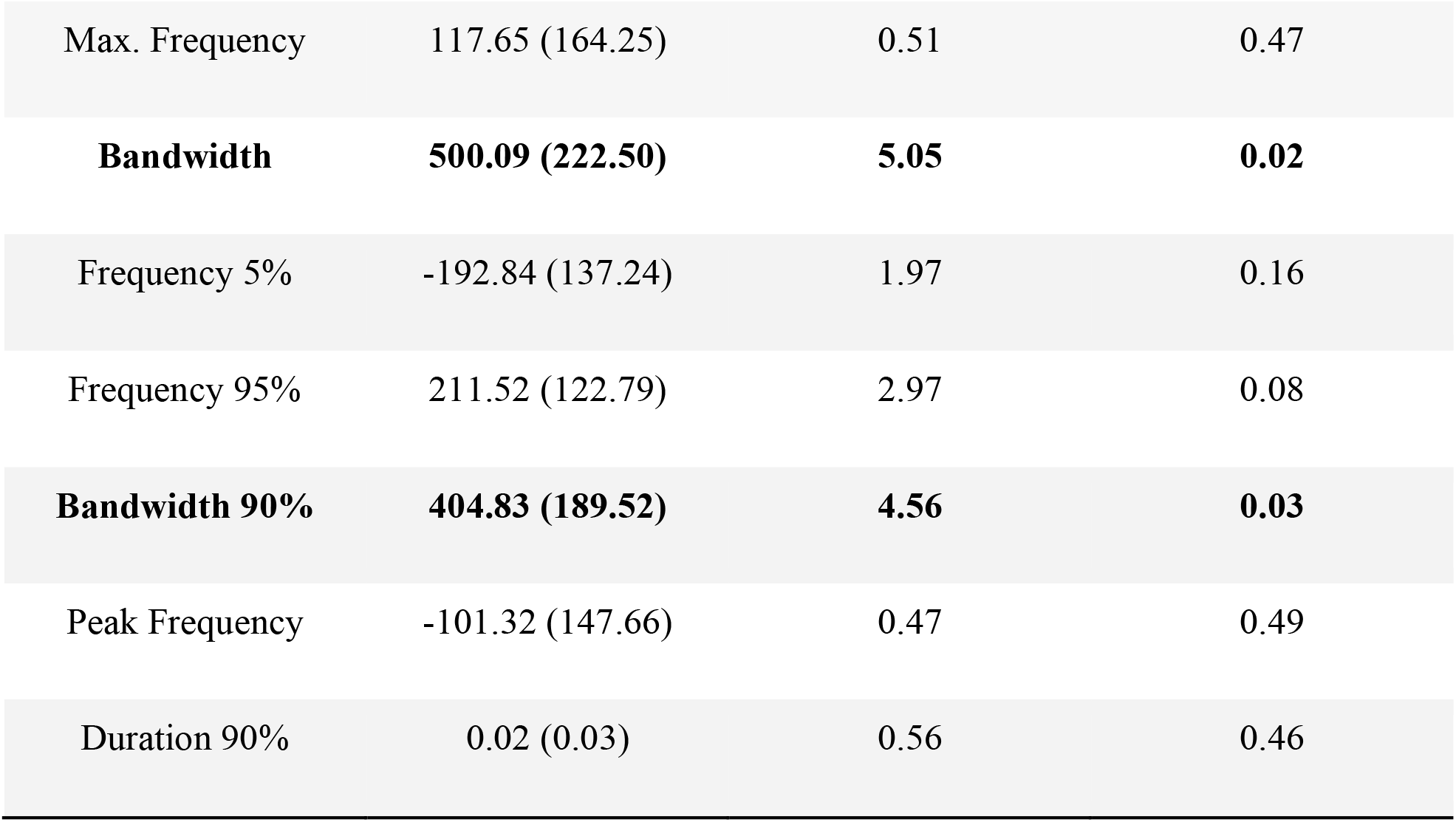
Linear mixed models for flourish phrases with the habitat type as the fixed factor (baseline = rural, n =343 for parameters measured with threshold method, for the rest n = 359)

Terminal Flourish Phrase
| | Coefficient (SE) | $\chi^2$ | P-value |
| --- | --- | --- | --- |
| **Min. Frequency** | **-385.29 (181.26)** | **4.53** | **0.03** |
| Max. Frequency | 117.65 (164.25) | 0.51 | 0.47 |
| **Bandwidth** | **500.09 (222.50)** | **5.05** | **0.02** |
| Frequency 5% | -192.84 (137.24) | 1.97 | 0.16 |
| Frequency 95% | 211.52 (122.79) | 2.97 | 0.08 |
| **Bandwidth 90%** | **404.83 (189.52)** | **4.56** | **0.03** |
| Peak Frequency | -101.32 (147.66) | 0.47 | 0.49 |
| Duration 90% | 0.02 (0.03) | 0.56 | 0.46 |

**Table 5.**
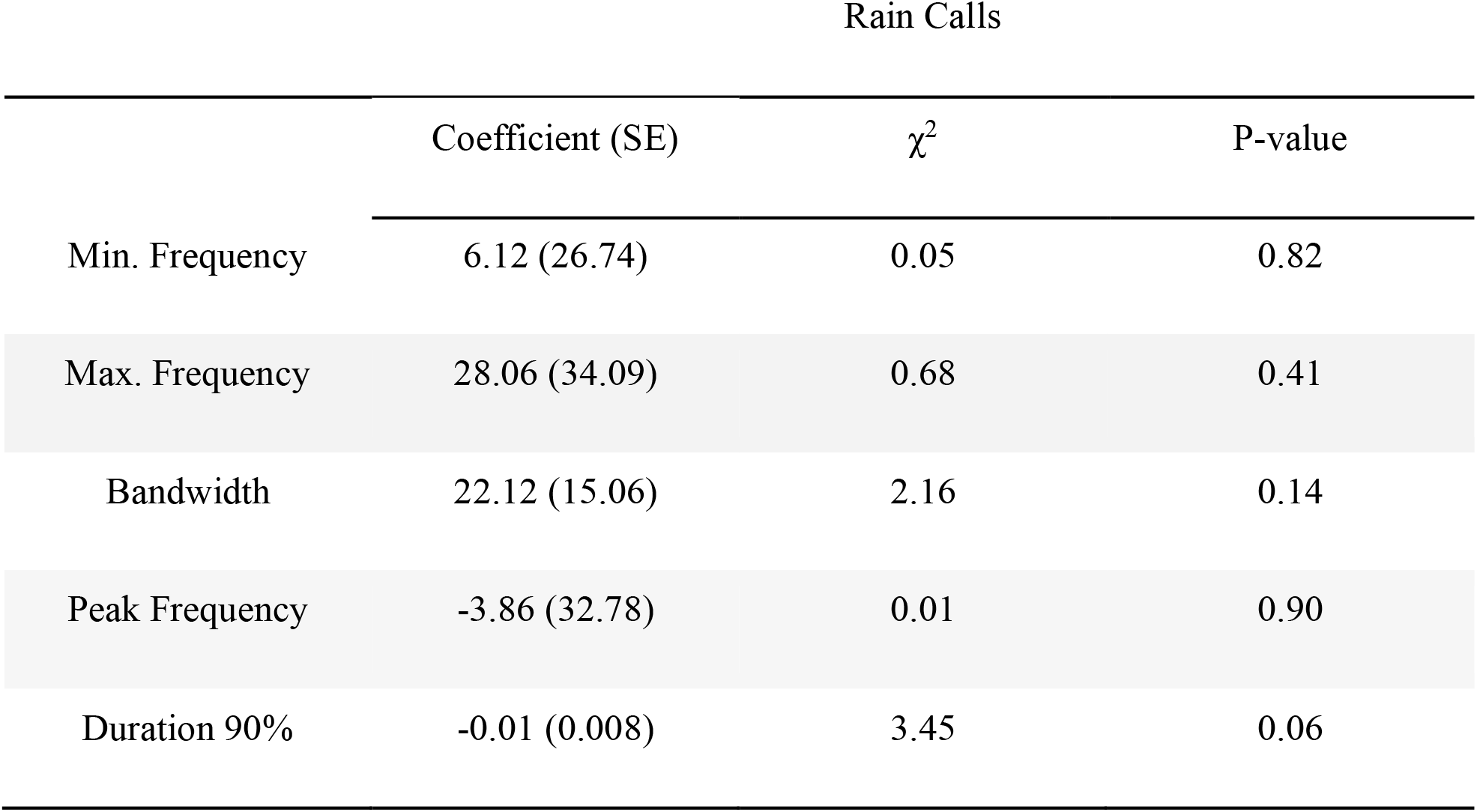
Linear mixed models for rain calls with the habitat type as the fixed factor (baseline = rural, n = 1570)

| Rain Calls |  |  |  |
| --- | --- | --- | --- |
| | Coefficient (SE) | $\chi^2$ | P-value |
| Min. Frequency | 6.12 (26.74) | 0.05 | 0.82 |
| Max. Frequency | 28.06 (34.09) | 0.68 | 0.41 |
| Bandwidth | 22.12 (15.06) | 2.16 | 0.14 |
| Peak Frequency | -3.86 (32.78) | 0.01 | 0.90 |
| Duration 90% | -0.01 (0.008) | 3.45 | 0.06 |

**Figure 3.**
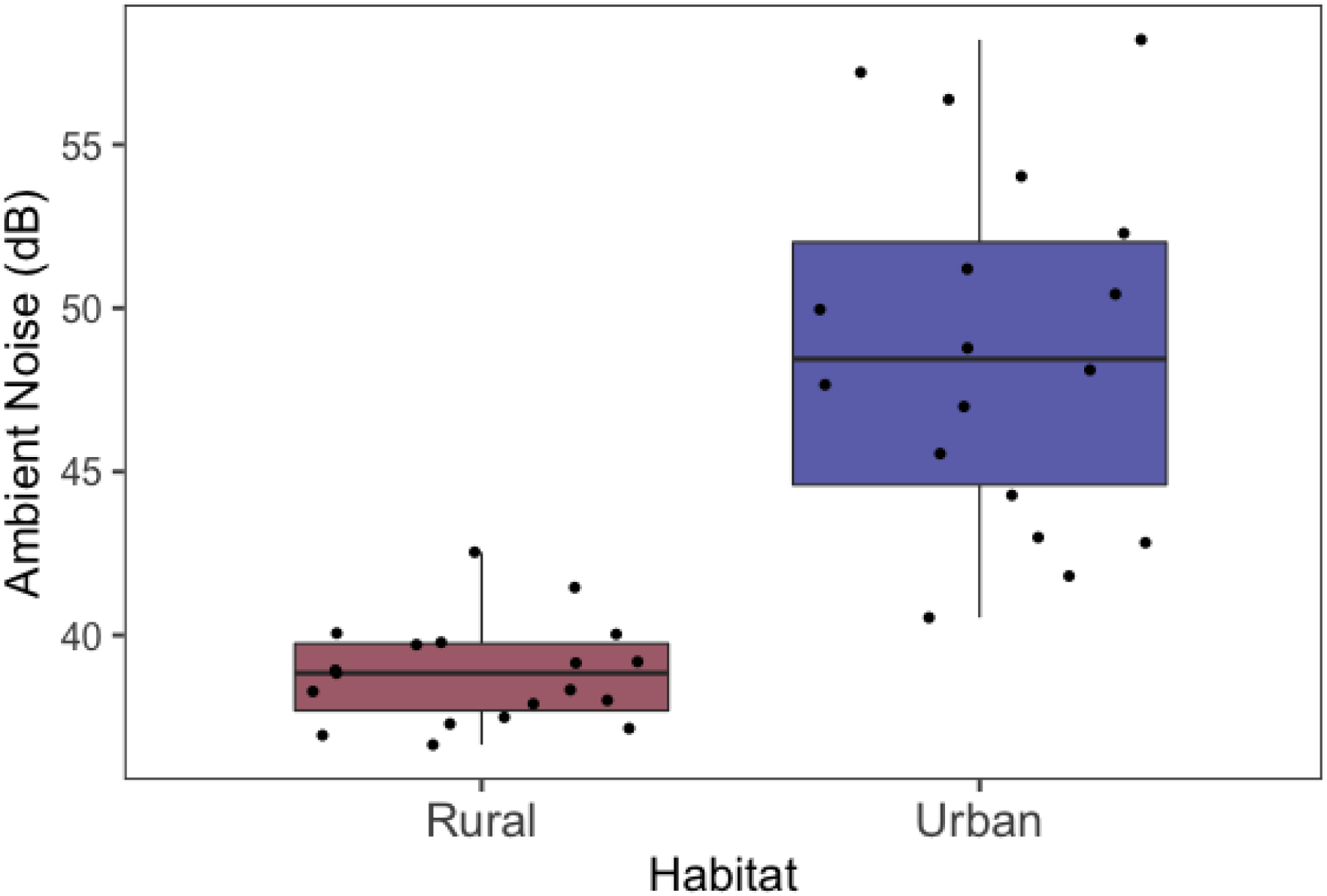
Boxplots of average ambient noise in chaffinch territories. Each data point indicates the ambient noise of a given day. Some territories were measured on two consecutive days.

**Figure 4.**
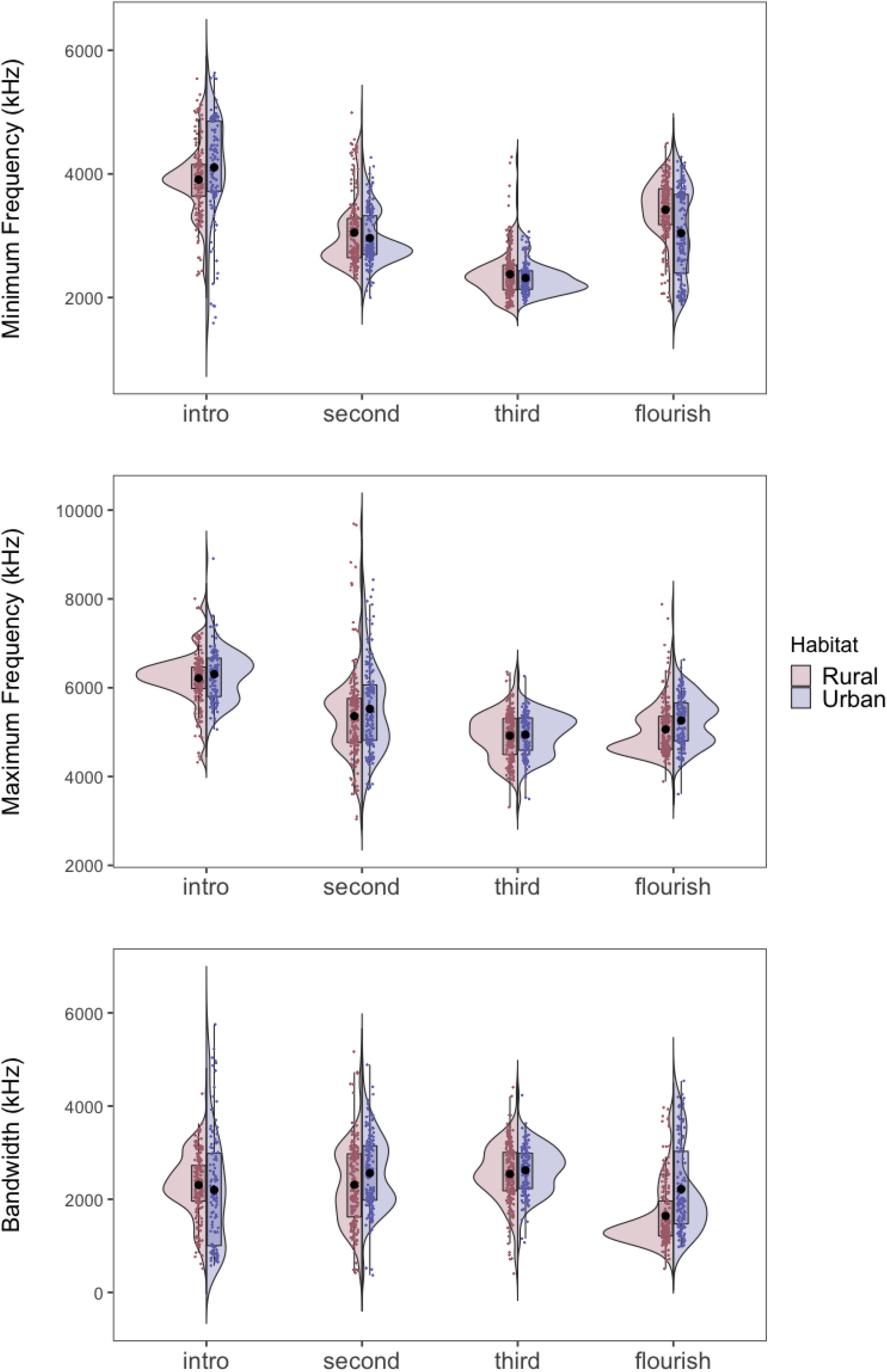
Split violin plots representing frequency parameter differences between songs recorded in urban and rural habitats. Black dots indicate means. From top to bottom: minimum frequencies, maximum frequencies, and bandwidths. Figure created using split violin plots from psychteachr/introdataviz (Nordmann et al., 2022).

## Discussion

Anthropogenic noise often affects the detectability of animal signals (Brumm & Slabbekoorn, 2005). In this study, we investigated whether chaffinches living in urban and rural habitats vocalize with different acoustic features which might increase the signal-to-noise ratio of their acoustic signals. Based on previous findings in other urban living songbirds (Brumm & Zollinger, 2013), we expected urban chaffinches to produce longer vocalizations with higher minimum frequencies and narrower bandwidths compared to rural chaffinches. Our results showed that while the rain calls did not differ at all, the effect of habitat varied depending on the phrase. Within the trilled phrases, there were no differences between urban and rural songs in any of the frequency or temporal characteristics we analyzed. Urban birds however, sang flourish notes which were significantly lower in minimum frequency, and higher in bandwidth than rural birds. The difference in bandwidth was about 500 Hz, which is likely a biologically meaningful difference (given the average bandwidth of the flourish is about 1900 Hz). We discuss these results in the context of the potentially different functions of the song phrases.

Several studies on species with trilled songs found that urban noise tends to lead to trill syllables with higher minimum frequencies in urban birds compared to songs from rural birds (Davidson et al., 2017; Job et al., 2016; Luther et al., 2016; Redondo et al., 2013). Instead we found that trill durations, rates, as well as frequency characteristics did not differ significantly between urban and rural males. This may be due to the fact that in our population, trills occupy generally higher frequencies with minimum frequencies well above 2000 Hz (see Figure 3). Given that most of the urban noise tends to be located below 1000 Hz, it may not interfere with effective transmission of the trill phrases amongst chaffinches. Nevertheless, other species with trilled phrases that occupy similar frequencies, such as white-crowned sparrows (*Zonotrichia leucophrys*), show increases of minimum frequencies in response to urban noise (Luther et al., 2016). The increases in minimum frequencies appear to be adaptive responses to improve transmission but also make the songs lower in trill performance which in turn reduces the effectiveness of the song as a signal to other males (Phillips & Derryberry, 2017).

While the chaffinch trills do not seem to vary in frequency and temporal characteristics between urban and rural habitats this lack of change may mean that urban songs are less effective signals in noisy urban habitats compared to rural habitats. Future studies determining responses to chaffinch songs from urban and rural habitats are required to test whether urban and rural songs draw differential responses in urban and rural habitats (e.g. Mockford & Marshall, 2009). Considering previous studies which suggested that chaffinch song phrases differ in their relative function with trills being more specialized for intrasexual communication and flourish phrases being more specialized for intersexual communication (Leitão & Riebel, 2003; Riebel & Slater, 1998), this comparison should include responses from both sexes.

### Why do urban flourishes have lower minimum frequencies and higher bandwidth?

The finding that the minimum frequency of flourish phrases decreases in the noisy urban habitats compared to rural habitats is contrary to our prediction. Most studies investigating the effect of anthropogenic noise on vocalizations found that minimum frequencies tend to shift upwards and often leading to a decrease in bandwidth (see reviews in Duquette et al., 2021; Roca et al., 2016). This increase in minimum frequencies along with narrower bandwidths may come about through an active change in the frequency of the songs to avoid masking by the noise (which tend to occupy lower frequencies), or it can be a side effect of a Lombard effect; the tendency to increase the amplitude of vocalizations in noise which may lead to an increase in frequency (Nemeth & Brumm, 2010; Zollinger et al., 2012). Both mechanisms are potentially adaptive in increasing the transmissibility of the vocalizations. Our finding of a lower minimum frequency and higher bandwidths of flourishes in the noisier urban habitats therefore is puzzling as on the face of it, it does the opposite.

It is worth noting, however, that a decrease in minimum frequency in urban habitats does not automatically mean lowered transmissibility of the signal. In a study on silvereyes (*Zosterops lateralis*), Potvin and colleagues (2014) found that alarm calls in urban habitats had lower frequency profiles (average, peak, and maximum frequencies) compared to calls in rural habitats. When they measured the detailed noise profiles of urban habitats, however, they found that the lowered frequency profiles of the alarm calls may actually increase the active space of the signal (the maximum distance at which the signal would be detected) in urban habitats relative to the frequency profiles of rural calls. It is possible that a similar process may operate here as well.

Findings showing the female directedness of flourish phrases raise the possibility that frequency differences only found in them are shaped by selection pressures posed by intersexual communication (Riebel & Slater, 1998). One speculation as to what those pressures might be in relation to the communication range required to interact with different sexes. Male-to-male communication usually occurs over short distances in territorial defense. On the other hand, to get mates or extra-pair copulations, signals should potentially travel over long distances. Depending on their colonization history, some species have much higher densities in urban habitats than in rural ones (Møller et al., 2012). We do not have such data for the chaffinches studied here, but urban areas with limited habitable territories may lead to denser populations compared to more continuous suitable habitats found in rural areas. Considering this and our personal observations, if the chaffinch population is sparser in rural habitats a signal with a larger active space for male-to-female communication would be adaptive. This could be achieved by having flourishes with narrow bandwidths as those kinds of signals have bigger active spaces in noise (Lohr et al., 2003).

Although chaffinches did not necessarily show adaptive changes in frequency and temporal characteristics of their signals in urban habitats, they may engage in other strategies to effectively communicate in noise (Brumm & Slabbekoorn, 2005). For instance, increasing the serial redundancy of the signal by repeating the same song type for longer bouts could be a strategy to adapt to noisy habitats used by chaffinches (Brumm & Slater, 2006; but see Deoniziak & Osiejuk, 2016). Moreover, other studies found that chaffinch song has higher minimum frequencies in noisy habitats (Brumm & Slater, 2006; Kislyakov & Ivanitskii, 2019). Note, however, that these latter studies manually estimated the minimum frequencies from spectrograms which is found to be an unreliable method (Brumm et al., 2017). Alternatively, chaffinches may also be increasing the amplitude of their song phrases in response to noise (Brumm & Zollinger, 2011). Our dataset lacks this measure as the distance between the recorder and the singing bird is not standardized, however, future research can investigate this possibility along with other strategies.

In summary, we found that despite the noisy conditions urban chaffinches did not have any of the measured strategies to overcome the masking caused by noise. On the other hand, the terminal flourish phrases had lower minimum frequencies and broader bandwidth in urban habitats, which possibly decreases the transmission efficiency of the phrase in noise. We speculate that population structure along with intersexual communication pressures could have shaped this difference.

## Ethics approval

This study did not require an ethics committee approval. All procedures used in this study follow the ASAB/ABS guidelines for the treatment of animals in behavioral research and teaching. Subjects were not captured or handled before, during or after any of the recordings. All recordings and playbacks lasted less than 5 minutes to minimize disturbance to the birds.

